# MINT: Predicting Enzyme-Cofactor Specificity by Multimodal Knowledge Transfer from Protein–Ligand Interactions

**DOI:** 10.64898/2026.10.08.757560

**Authors:** Yilin Ye, Xinkang Li, Hongliang Duan, Yingfeng An, Jingjing Guo

## Abstract

Protein-cofactor selectivity prediction is important for enzyme annotation, metabolic engineering, and biocatalyst discovery, but it remains challenging because experimentally supported associations are limited and unevenly distributed, and because biologically relevant differences can be subtler than those captured by general protein-ligand interaction models. Here, we present MINT (Multimodal Interaction Network with Teacher supervision), a multimodal framework for cofactor specificity prediction. MINT combines protein sequence features with cofactor sequence and graph representations and is trained with a gradual adaptation strategy that transfers useful information from a pretrained protein-ligand interaction model to the cofactor prediction task. On a fixed balanced held-out test set, MINT achieved the best overall performance among all compared methods (AUC = 0.932), and this ranking was reproduced in an independent family-stratified evaluation at the natural positive rate (pooled AUC = 0.939). Further analyses suggest that the pretrained model provides transferable structural information, while the gradual training scheme helps adapt that information to cofactor specificity. In a case study of the chemically near-identical NAD/NADP pair, MINT recovered 95.0-98.6% of true binders and resolved a single-phosphate-level distinction with direct relevance to engineering enzyme cofactor specificity. These results show that gradual transfer learning is an effective approach to cofactor specificity prediction when annotation is limited.

**TOC:** 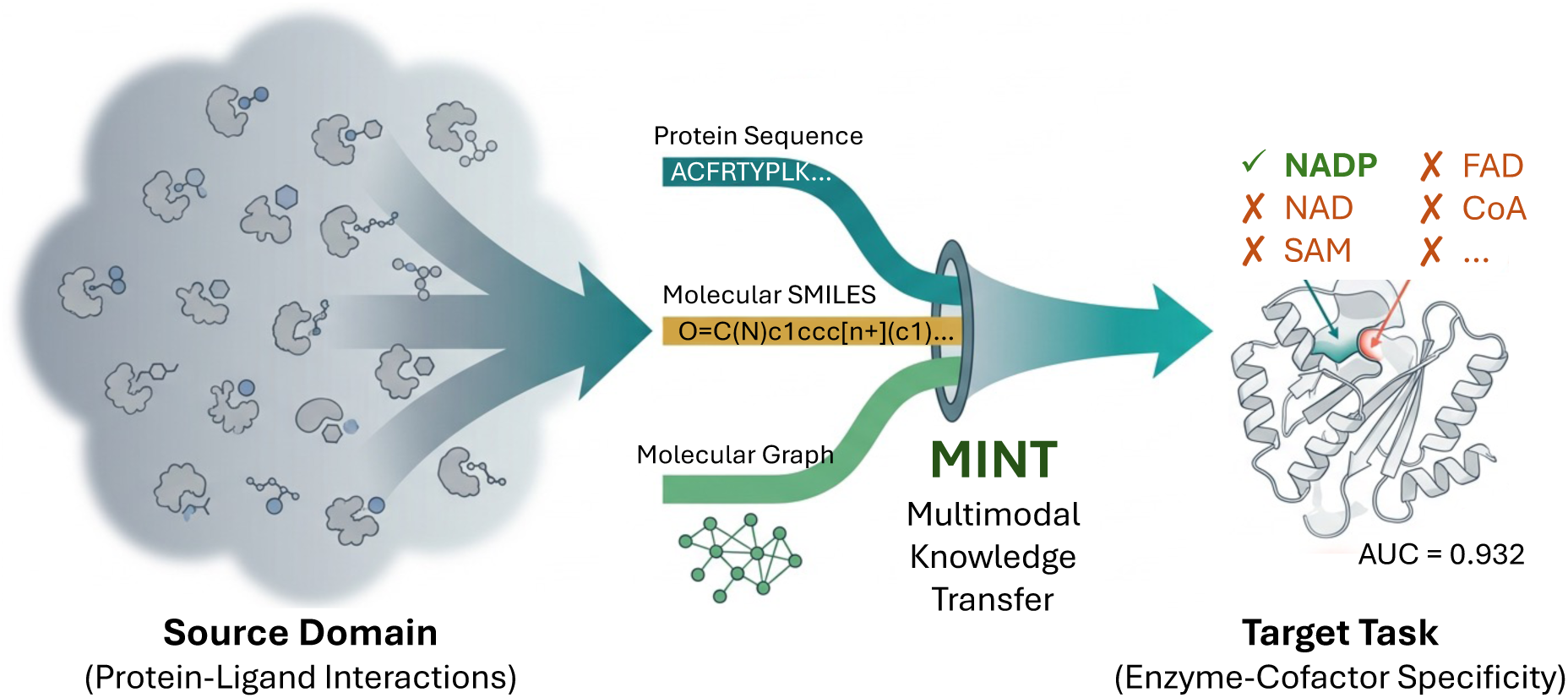

## INTRODUCTION

Accurate prediction of protein-cofactor selectivity is increasingly important for enzyme annotation, metabolic engineering, and biocatalyst discovery, because protein-cofactor associations are central to catalysis, metabolic regulation, and cellular homeostasis [1], [2]. We use the term cofactor in a broad sense throughout, including organic coenzymes as well as molecules such as heme that are not strictly classified as coenzymes [3]. Cofactors participate directly in essential chemical processes, including electron transfer and group transfer, so differences in cofactor usage are closely related to catalytic behavior and to the organization and function of metabolic networks [4], [5]. At the same time, the need for reliable computational prediction has grown because experimental characterization has not kept pace with the rapid increase in unannotated proteins [6]. The main difficulty is therefore not the importance of the problem, but the limited and uneven availability of reliable labels. Relative to the vast combinatorial space of proteins and biologically active small molecules, experimentally validated protein-cofactor associations remain scarce, and the available examples are distributed unevenly across cofactor classes and protein families [7]. As a result, models trained directly on task-specific data are vulnerable to overfitting, lineage bias, and weak generalization, even when they perform well on narrow benchmarks [8].

This challenge also helps explain why conventional approaches are often insufficient. Earlier studies on related biochemical prediction tasks have typically relied on curated supervision, using either manually designed features or relatively simple neural models [9], [10]. These methods can perform well when annotations are abundant and closely aligned with the target task, but protein-cofactor selectivity requires finer discrimination. Although this task belongs to the broader area of molecular recognition, direct supervision for it is much sparser than for general protein-ligand interaction prediction [11], [12], [13]. The central difficulty is that the task depends on subtle biological distinctions, yet provides only limited evidence from which those distinctions can be learned reliably.

Against this background, recent advances in machine learning make transfer learning a practical alternative, because knowledge learned from large related biochemical datasets can be adapted to narrower downstream tasks [14], [15], [16]. Knowledge distillation further strengthens this idea by transferring predictive structure from a stronger or more general model to a task-specific model through soft targets, allowing the downstream learner to benefit not only from scarce direct labels but also from broader regularities in molecular interactions [17].

From this perspective, cofactor prediction is better understood not as an isolated problem, but as a target task that partially overlaps with a broader source domain. General protein-ligand interaction and protein-cofactor recognition share common physicochemical principles, but they are not the same objective [18]. A model trained only for broad interaction prediction may learn useful binding patterns without being well tuned to cofactor selectivity, whereas a model trained only on the small cofactor dataset may align more closely with the final objective but lack the prior knowledge needed for robust learning. The methodological challenge is therefore to retain transferable knowledge while adapting the model to a cofactor-specific objective. This issue is closely tied to the transition between tasks: if optimization shifts too abruptly from the source task to the target task, knowledge acquired in the broader domain may be lost, especially when downstream annotations are sparse and noisy [19], [20]. An effective framework should therefore support gradual specialization, so that generally useful recognition patterns are preserved while the model becomes better suited to finer cofactor discrimination.

These considerations also have direct implications for model design. Because limited data increase the risk of overfitting, earlier studies have often favored relatively simple architectures to maintain training stability [21], [22], [23]. Although this choice is reasonable, it can limit the model’s ability to capture more complex determinants of biochemical recognition. If a model first receives large-scale indirect supervision, however, a more expressive architecture becomes easier to train and can begin downstream adaptation from a more informative parameter state. The performance achievable in cofactor prediction therefore depends not only on the final objective but also on the training path used to reach it.

Motivated by this view, this study proposes MINT (Multimodal Interaction Network with Teacher supervision). Protein-cofactor selectivity is formulated as a pairwise prediction task in which each protein-cofactor pair is labeled as positive or negative according to available experimental evidence. The framework first introduces large-scale soft supervision from a broader interaction domain, allowing the target model to learn general molecular recognition patterns before focusing more directly on the cofactor-related objective. It then applies distillation-based initialization followed by downstream adaptation, improving both parameter initialization and the practical value obtained from limited supervision. To narrow the gap between source and target tasks, we further design a progressive adaptation strategy that shifts optimization step by step from general interaction modeling to cofactor selectivity prediction, helping preserve transferable knowledge and support more stable specialization. Building on this design, we also explore a more expressive student architecture that can make fuller use of the richer supervisory signal provided by transfer learning and distillation. Overall, the proposed framework combines indirect large-scale supervision, progressive cross-task adaptation, and greater model expressiveness to improve prediction under limited annotation and strengthen enzyme-related functional inference.

## MATERIALS AND METHODS

### Datasets

#### Cofactor category integration

Based on the study by Mukhopadhyay et al., we obtained a JSON file to systematically record annotations for 365 unique cofactors, each identified by its PDB Chemical Component ID [7]. For downstream analysis, we built two complementary cofactor classification frameworks by combining chemical standardization with functional consolidation.

We first extracted all three-character cofactor codes from the input dataset and normalized them into unique entries to reduce duplication in the annotations. Each normalized code was then mapped to the corresponding record in the Chemical Component Dictionary of the RCSB Protein Data Bank, which provided compound names, classifications, alternative names, and relevant chemical properties [24]. We standardized molecular structures with RDKit by parsing each cofactor structure and converting it to a canonical SMILES string, so that equivalent structures submitted under different codes or tautomeric forms were merged into a single chemical identity [25].

We then resolved the remaining redundancy at the molecular level using a hierarchical matching strategy, with InChIKey as the primary descriptor and InChI or canonical SMILES used when they provided a more reliable representation [26], [27]. Entries sharing the same chemical descriptor were grouped into an identity cluster, and one representative code was retained for consistency. We next added a second integration step to remove distinctions that were chemically annotated but not biologically informative, especially for cofactors that played the same biochemical role despite differences in redox state or naming conventions. These chemically distinct but functionally equivalent entries were merged into cofactor families [28]. This procedure produced both a chemically non-redundant cofactor set and a functionally integrated family set [29]. After consolidation, 265 cofactors were retained.

#### Compilation of protein cofactor preference data

To build a reliable dataset of protein cofactor preferences, we followed the data processing strategy of Ye et al. and integrated information from multiple databases to improve both coverage and consistency [30]. The resulting annotations were merged into 11,477 protein-cofactor pair records and then deduplicated using the procedure described by Ye et al.: proteins were clustered by pairwise sequence identity at a threshold of 0.4, and records within each cluster were consolidated, yielding the final dataset used for modeling (Figure 1A). The final dataset contains 4,262 experimentally supported positive pairs spanning 265 cofactors and 3,912 proteins. By exhaustively combining the retained proteins and cofactors, we generated 1,036,680 protein-cofactor pairs in total. Among these, the 4,262 experimentally supported pairs were labeled as positive samples, whereas the remaining 1,032,418 pairs were treated as negative samples (Table 1). The distribution of positive samples across cofactor families is highly concentrated: the ten most abundant families account for most positive samples, whereas eleven families contain fewer than 50 positives each (Figure 1B).

**Figure 1.**
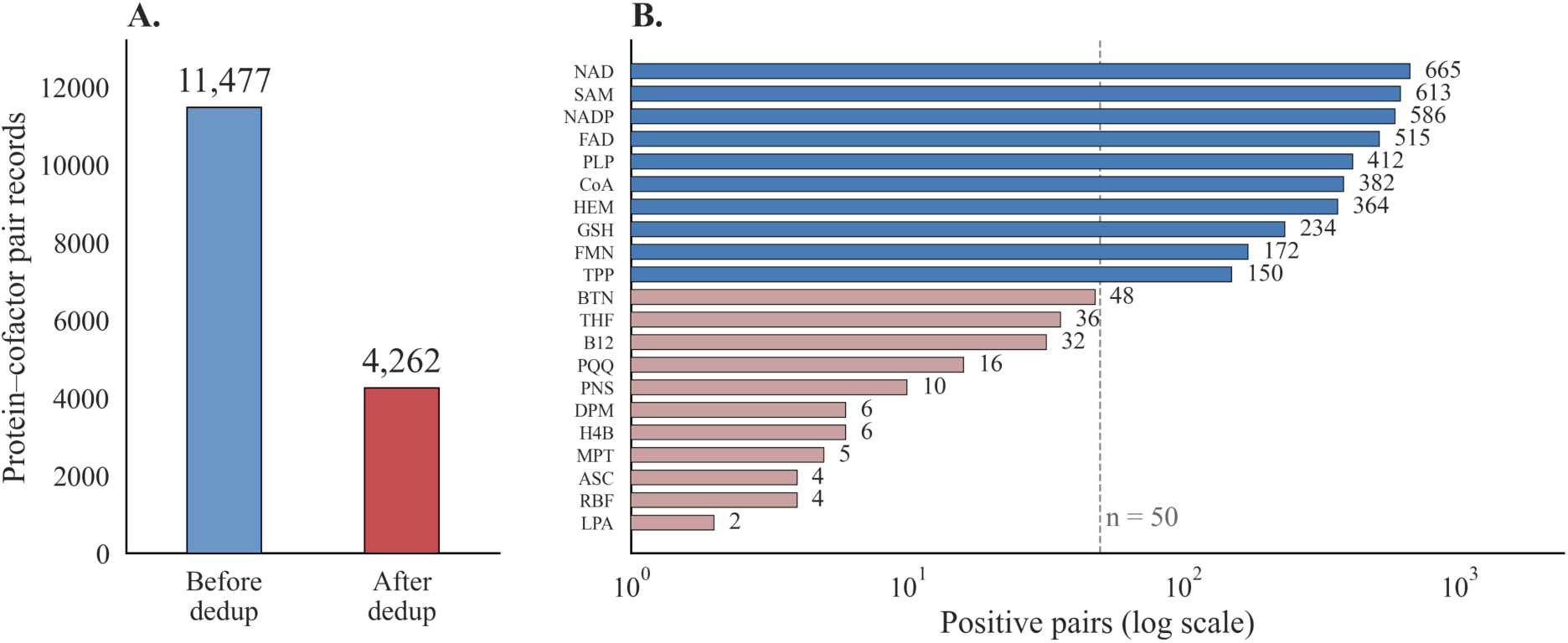
Dataset curation and family-level composition. (A) Record-level deduplication workflow. Annotations integrated from multiple databases were merged into 11,477 protein-cofactor pair records and deduplicated using the procedure described by Ye et al., yielding the 4,262 positive pairs used for modeling. (B) Number of positive pairs in each cofactor family (log scale; dashed line, *n*=50). Abbreviations: NAD(P), nicotinamide adenine dinucleotide (phosphate); SAM, S-adenosylmethionine; FAD, flavin adenine dinucleotide; PLP, pyridoxal 5′-phosphate; CoA, coenzyme A; HEM, heme; GSH, glutathione; FMN, flavin mononucleotide; TPP, thiamine pyrophosphate; BTN, biotin; THF, tetrahydrofolate; B12, cobalamin; PQQ, pyrroloquinoline quinone; PNS, phosphopantetheine; DPM, dipyrromethane; H4B, tetrahydrobiopterin; MPT, molybdopterin; ASC, ascorbate; RBF, riboflavin; LPA, lipoate.

**Table 1.** Statistics of the curated dataset and the evaluation splits.

| Item | Value | Note |
| --- | --- | --- |
| Proteins | 3,912 | After deduplication following the procedure of Ye et al. |
| Cofactors | 265 | After canonical SMILES standardization and family consolidation |
| Enzyme-cofactor pairs | 1,036,680 | Exhaustive pairing |
| Positive pairs | 4,262 | Experimentally supported; natural positive rate 0.41% |
| Held-out test set | 1,000 positive + 1,000 negative | Fixed across all configurations |
| Cross-validation data | Remaining pairs | Five-fold; five models evaluated on the same test set |

### Data splitting

For the standard evaluation setting, we randomly sampled 1,000 positive pairs and 1,000 negative pairs to form a fixed held-out test set (Table 1). The remaining data were used for five-fold cross-validation. In each fold, a separate model was trained, with the validation set used for hyperparameter tuning and model selection, and all five models were then evaluated on the same fixed test set. Performance is reported as the mean ± standard deviation across the five models. This design provides a clear separation between model development and final testing and allows fair comparison across model variants.

Because standard cross-validation on exhaustively paired data can produce overly optimistic estimates, we also constructed a family-stratified five-fold split. Cofactors were grouped into families according to the chemical and functional criteria described above, and each family-by-label group was divided into five parts that rotated through training and testing. This ensured that every sample was held out exactly once and that performance could be aggregated over the full dataset at its natural positive rate (0.41%). Hyperparameters were inherited from the standard setting without further tuning. Results from this split are reported separately from those of the standard setting and are not directly comparable to the balanced-test results.

### Model architecture

#### Teacher–student learning framework

Our approach for protein-cofactor selectivity prediction is based on a teacher-student framework that emphasizes knowledge transfer rather than relying only on hard labels. In this framework, a pretrained teacher model assigns soft probabilistic scores to each protein-cofactor pair, allowing the student to learn finer interaction patterns and latent preferences beyond binary annotations (Figure 2A).

**Figure 2.**
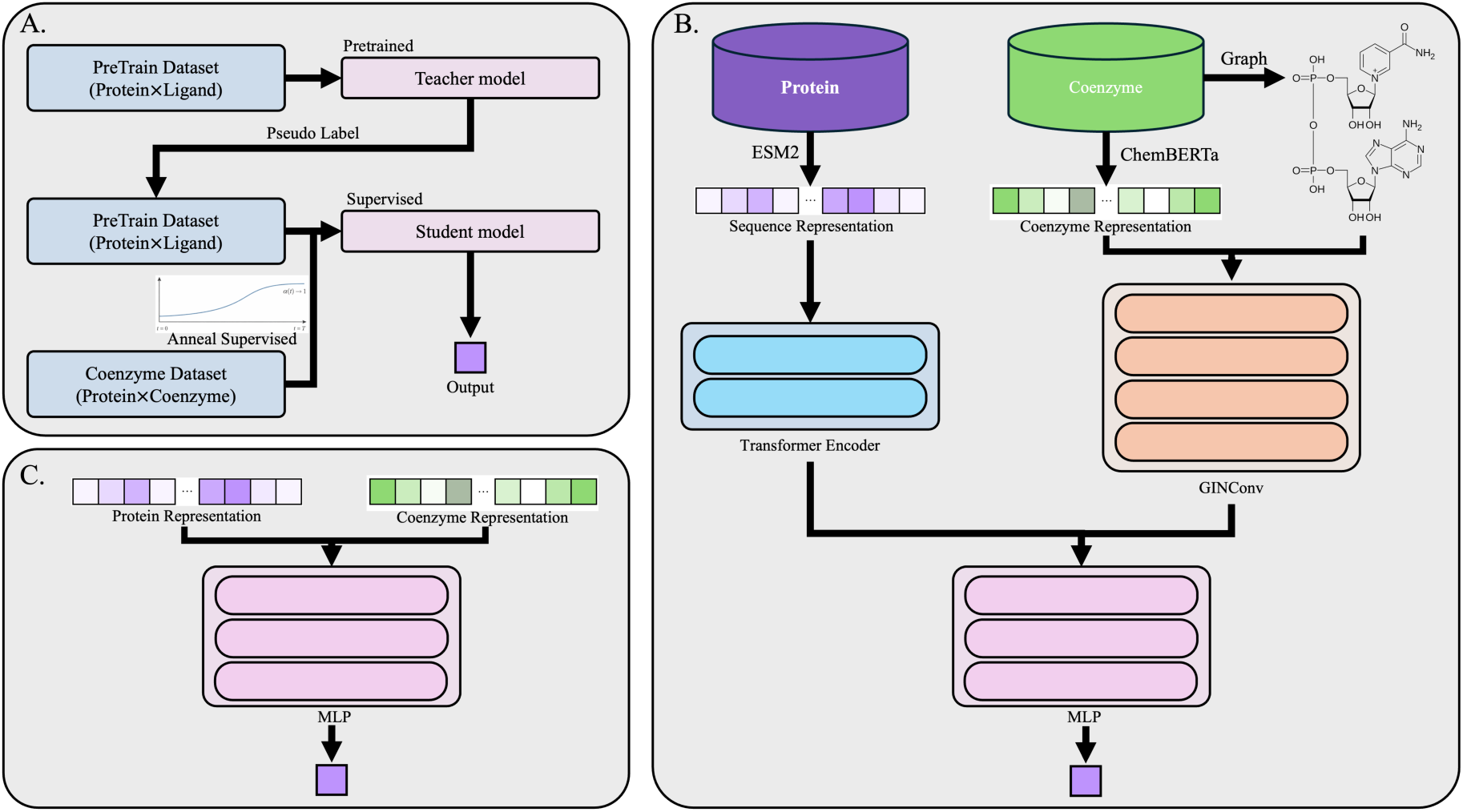
Overview of the teacher-student training strategy and the MINT framework for general cofactor specificity prediction. (A) Schematic of the teacher-student training procedure. (B) Workflow of the proposed MINT framework. (C) MLP-based baseline for cofactor specificity prediction.

We used PLXFPred as the initial teacher model. PLXFPred is a multimodal deep learning framework developed for protein-ligand interaction classification and binding affinity prediction [31]. It uses separate encoders to extract graph-level and sequence-level representations for both proteins and ligands. In the protein branch, physicochemical, positional, and evolutionary features are integrated together through a two-layer GATv2-based graph encoder and a BiLSTM-based sequence encoder. In the ligand branch, PLXFPred uses a three-stream graph encoder consisting of a molecular graph branch, a clique or tree-decomposition branch, and a ChemBERTa-based semantic branch, whose outputs are fused by self-attention [32]. Graph and sequence features are then integrated through a bidirectional cross-attention module, in which graph representations attend to sequence representations and vice versa, followed by gated fusion. To improve representation learning, the model adopts a hierarchical multimodal fusion strategy that combines raw graph features, early interaction features, and cross-modal fused features through a Transformer-based fusion layer with residual connections. The final fused representation is passed to parallel prediction heads for affinity regression, activity classification, and domain-adversarial generalization.

The teacher model was first used to predict binding probabilities for the generated training instances, producing soft labels. Compared with hard class annotations, these soft labels retain more nuanced information about inter-class similarity and prediction uncertainty. The student model was first initialized by pretraining on these soft labels and was then further optimized for the downstream task. This strategy allowed the model to learn from a broader supervisory signal during pretraining while reducing the effect of the severe shortage of labeled examples in some classes of the original dataset.

#### Distillation and annealed optimization

The student is a multimodal predictor that integrates protein sequence information, cofactor SMILES representations, and graph-based topological features of cofactors within a unified architecture (Figure 2B). By combining these complementary modalities under the teacher’s guidance, the student learns general principles of enzyme-cofactor recognition from the teacher and adapts them to the specific task of cofactor selectivity prediction. For each enzyme-cofactor pair, the ground-truth interaction label is denoted by *y*, the soft probability produced by the teacher by *p*, and the student outputs a scalar logit *z*, which is converted to a predicted probability by the logistic sigmoid function *σ*.

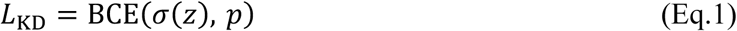

To guide early optimization, training first introduces a knowledge distillation objective (Eq.1), which encourages the student to match the teacher’s output distribution more closely and helps stabilize the initial learning process. Rather than relying only on fixed class assignments, the model can learn from the teacher’s soft labels, which capture relational structure as well as confidence information. This richer signal provides a more informative starting point for the subsequent stage of task-specific optimization.

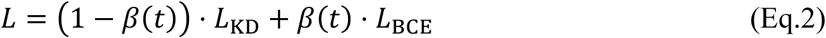

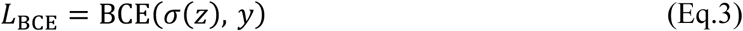

After the distillation stage, the student network is further optimized using a hybrid supervision scheme that combines hard targets with guidance from the teacher model (Eq.2), where Eq.3 is the binary cross-entropy with respect to the ground-truth labels. Instead of keeping the mixing coefficient fixed throughout training, we vary this epoch-dependent weight to adjust the balance between distillation and supervised learning over time, following a cosine annealing schedule:

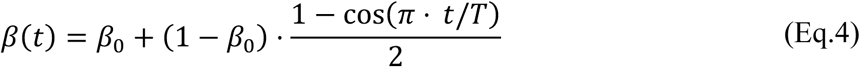

where *t* denotes the current training epoch and *T* the total number of fine-tuning epochs. The weight on hard labels therefore increases from 0.1 at the start of fine-tuning to 1.0 at the end. Early training is driven mainly by supervision from the teacher, whereas later stages place increasing emphasis on the ground truth, allowing the student to move smoothly from teacher imitation to task-specific discrimination. In the configuration without annealing (w/o A), the second stage optimizes only the hard-label objective (Eq.3), that is, *β* is fixed at 1 rather than scheduled.

#### Student architecture

To improve interaction prediction beyond what any single modality can offer, the student architecture adopts a multimodal design that combines complementary information from enzymes and cofactors. The workflow begins with two dedicated encoders that learn enzyme and cofactor representations separately and then integrate them for downstream prediction. In the protein branch, the enzyme sequence is first encoded with the pretrained ESM2 model (esm2_t33_650M_UR50D), which serves as a frozen feature extractor [33]. Per-residue embeddings of 1,280 dimensions are generated offline, and the model parameters are not updated during training. The resulting sequence representation is then passed through a two-layer Transformer encoder (four attention heads, hidden dimension 256, feed-forward dimension 1,024) to capture contextual dependencies and produce a refined enzyme representation.

For the cofactor branch, we used a pretrained ChemBERTa model to generate context-aware 768-dimensional embeddings from SMILES sequences, together with a 2,048-bit Morgan fingerprint; both were projected into a shared 256-dimensional space. The same cofactor was also represented as a molecular graph and processed by four GINConv layers (256 dimensions) to preserve atom-level relationships and global structural topology [34]. Before message passing in the GINConv layers, the SMILES-derived features were aligned with graph-based atom features and incorporated into the node representations as either primary or auxiliary information. This allows the encoder to capture sequential chemical semantics and topological molecular structure within a unified representation. The final cofactor embedding was obtained by aggregating features across all nodes.

After the protein and cofactor were encoded separately, their embeddings were integrated through a bidirectional multi-head cross-attention module (four heads, 256 dimensions), in which the protein representation attends to the cofactor representation and the cofactor representation attends to the protein representation, yielding a joint representation of the enzyme-cofactor pair [35]. A feedforward neural network (1,536 → 512 → 1) was then used to compute the predictive logit from this integrated encoding, and the final score was converted into a probability by the sigmoid function. For a more rigorous comparative evaluation, we also constructed an MLP-based baseline following the general design of INSIGHT. In this baseline, the enzyme and cofactor were first encoded into their respective representations and then passed to a single-hidden-layer MLP (hidden dimension 512) for final prediction (Figure 2C).

#### Training details and model selection

All models were optimized with AdamW (learning rate 3×10^−4^, weight decay 10^−2^) using a batch size of 32 and a fixed random seed [36]. Training was conducted in two stages: a distillation (pretraining) stage on teacher soft labels, followed by a fine-tuning stage under the annealed hybrid objective. Each stage was trained for up to 20 epochs with early stopping and terminated if the validation AUC did not improve for 5 consecutive epochs. For each configuration, the checkpoint with the highest validation AUC was retained. Hyperparameters were selected by grid search based on cross-validation performance on the validation folds.

## RESULTS

### Model architecture and configuration overview

The details of the proposed framework and training procedure are described in Materials and Methods (Figure 2). Briefly, the training is conducted in two stages: the student first learns to reproduce the teacher’s soft scores through distillation (D), and the objective is then gradually shifted toward the hard experimental labels under a cosine annealing schedule (A).

Using the two backbones, MINT and an MLP baseline following the design of INSIGHT, we defined six configurations that differ only in training strategy (Table 2): the full pipeline (D+A), distillation without annealing (w/o A), and supervised training without distillation (w/o D). The teacher, which is not trained within this framework, is included as an additional reference in all evaluations.

**Table 2.** Configurations of the compared models.

| Backbone | Configuration | Distillation (D) | Annealing (A) |
| --- | --- | --- | --- |
| MINT | D+A (full) | ✓ | ✓ |
|  | w/o A | ✓ | ✗ |
|  | w/o D (supervised) | ✗ | ✗ |
| MLP | D+A (full) | ✓ | ✓ |
|  | w/o A | ✓ | ✗ |
|  | w/o D (supervised) | ✗ | ✗ |

### Overall performance under cross-validation

We first evaluated all models under the standard cross-validation setting. The curated dataset contained 1,036,680 exhaustively paired enzyme-cofactor pairs derived from 3,912 proteins and 265 cofactors, of which 4,262 were supported by experimental evidence (Materials and Methods). For each of the two student backbones, we compared the three training configurations defined in Table 2. Before training, we reserved a fixed held-out test set of 2,000 balanced pairs (1,000 positive and 1,000 negative). For each configuration, five models were trained via five-fold cross-validation on the remaining data, with each model evaluated on the same test set. The means and standard deviations of the main indicators across the five models are summarized in Table 3 and Figure 3.

**Figure 3.**
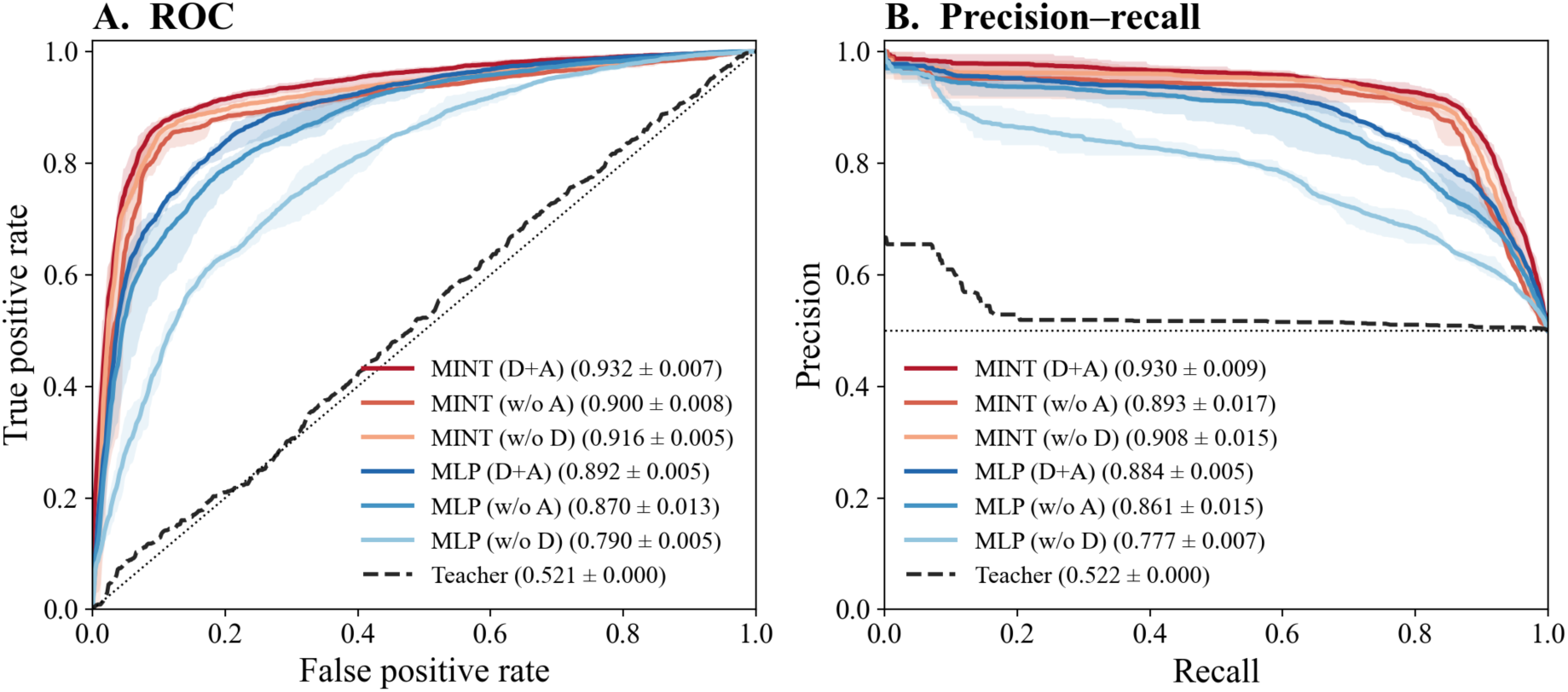
ROC (A) and precision-recall (B) curves on the held-out test set for all six configurations and the teacher. Shaded bands indicate the range across the five cross-validation models, and solid lines show the mean curve. The legend reports the mean ± SD of AUC (A) and average precision (B) across models. The teacher (dashed) is independent of the training procedure and is therefore identical across folds, and its band collapses to the curve itself.

**Table 3.** Overall performance on the held-out test set (mean ± SD across the five cross-validation models).

| Backbone | Configuration | AUC | AUPR | ACC |
| --- | --- | --- | --- | --- |
| MINT | D+A (full) | **0.9320 ± 0.0065** | **0.9298 ± 0.0092** | **0.8816 ± 0.0049** |
| | w/o A | 0.9002 ± 0.0080 | 0.8931 ± 0.0170 | 0.8657 ± 0.0048 |
| | w/o D (supervised) | 0.9159 ± 0.0053 | 0.9079 ± 0.0151 | 0.8654 ± 0.0114 |
| MLP | D+A (full) | **0.8920 ± 0.0055** | **0.8839 ± 0.0050** | **0.8126 ± 0.0080** |
| | w/o A | 0.8697 ± 0.0128 | 0.8611 ± 0.0150 | 0.7929 ± 0.0085 |
| | w/o D (supervised) | 0.7903 ± 0.0049 | 0.7775 ± 0.0070 | 0.7115 ± 0.0135 |
| Teacher | - | 0.5211 | 0.5221 | 0.5095 |

As can be seen, MINT with the full pipeline achieved the best overall performance (AUC = 0.932, AUPR = 0.930, ACC = 0.882), outperforming all ablated variants and the MLP baseline on every metric. The two MINT ablations remained competitive (w/o D: 0.916; w/o A: 0.900), with their rankings relative to the full model unchanged. The MLP baseline also benefited substantially from teacher-guided training (D+A: 0.892 vs. w/o D: 0.790; difference: 0.102), although it still underperformed all MINT configurations.

In contrast, the teacher model itself performed near chance on this task, achieving an AUC of only 0.5211 against the true labels on the same held-out test set. This indicates that general protein-ligand interaction scores alone are not sufficient to rank cofactor preferences effectively. Nevertheless, both student backbones substantially outperformed the teacher across all configurations that incorporated its soft labels, suggesting that the teacher’s value lies in the supervisory signal it provides rather than in its direct predictions.

### Distillation and annealing: ablation and pseudo-teacher controls

To understand where the benefit of teacher-guided learning comes from, we extended the comparison in two directions. We first constructed pseudo-teacher controls in which the soft labels were replaced by either random values or shuffled versions of the real teacher outputs. The shuffled labels preserve the exact marginal score distribution but break the correspondence between teacher scores and enzyme-cofactor pairs (Table 4, Figure 4). We then ablated each component of the training strategy on both backbones and compared the full pipeline with its variants across the five cross-validation models (Figure 5). All configurations used the same data splits and training protocol; only the soft-label source changed.

**Figure 4.**
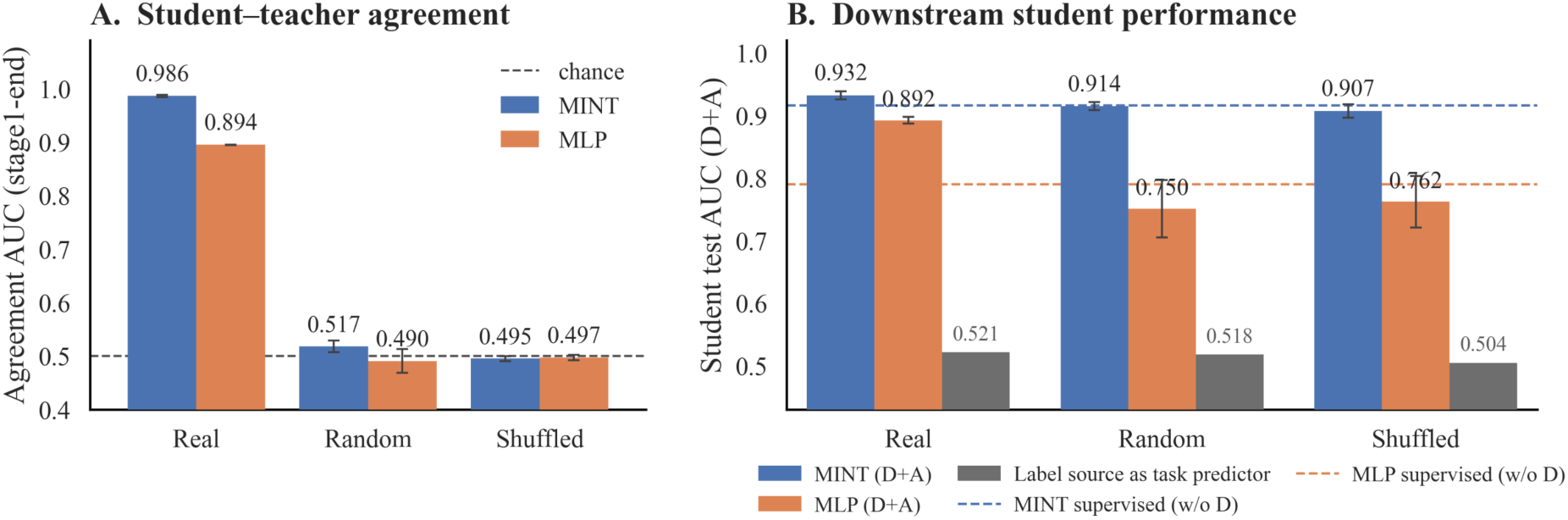
Task alignment and information content of the teacher signal are orthogonal. (A) Student-teacher agreement at the end of distillation (agreement AUC on the held-out test set, mean ± SD across five models). Only the real teacher is reproduced faithfully (0.986 for MINT, 0.895 for MLP); random and shuffled labels remain at chance level (dashed line). (B) Final student test AUC after the full pipeline was trained on each label source (mean ± SD, five-fold cross-validation). Grey bars show the AUC of each label source itself against the ground-truth labels on the same held-out test set (real 0.521, random 0.518, shuffled 0.504, all near chance). Dashed lines indicate the supervised references trained without distillation. Label sources that are indistinguishable as task predictors lead to sharply different downstream outcomes because only the real teacher provides structured, transferable knowledge.

**Figure 5.**
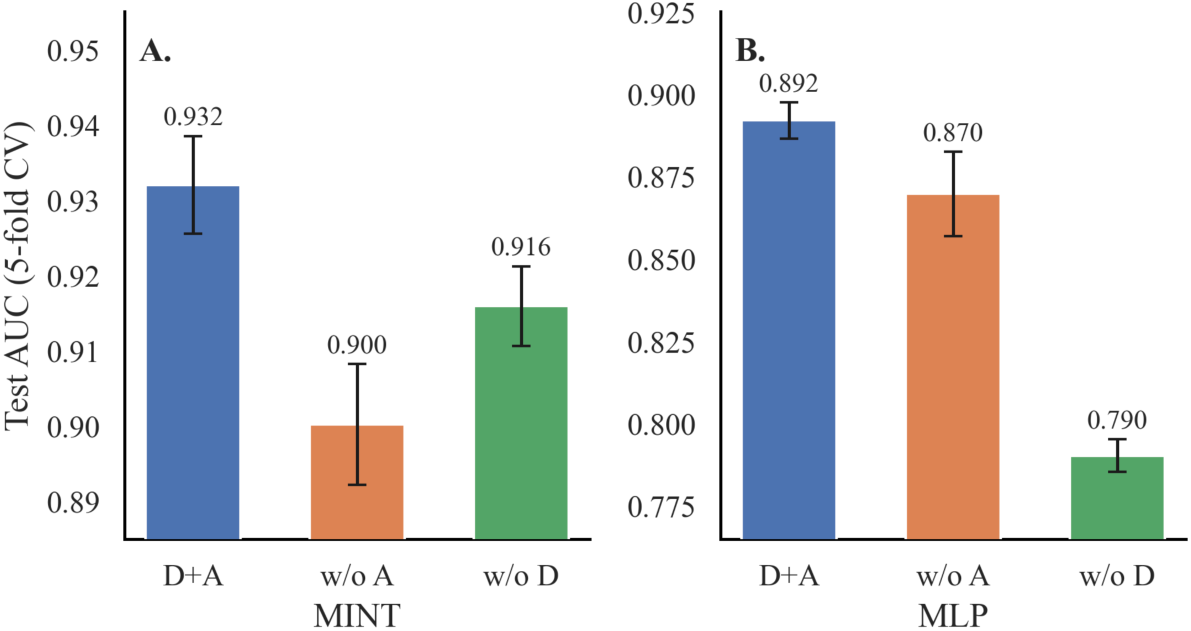
Contribution of distillation and annealing on the two backbones. Mean test AUC (± SD across the five cross-validation models on the held-out test set) for the full pipeline (D+A), without annealing (w/o A), and without distillation (w/o D).

**Table 4.** Distillation with real and pseudo-teacher soft labels (mean ± SD, five models, held-out test set).

| Backbone | Soft-label source | Configuration | AUC | AUPR |
| --- | --- | --- | --- | --- |
| MINT | Real teacher | D+A | **0.9320 ± 0.0065** | **0.9298 ± 0.0092** |
| MINT | Random | D+A | 0.9144 ± 0.0063 | 0.9137 ± 0.0121 |
| MINT | Shuffled teacher | D+A | 0.9069 ± 0.0106 | 0.9094 ± 0.0147 |
| MINT | Random | w/o A | 0.9028 ± 0.0063 | 0.8974 ± 0.0104 |
| MINT | - | w/o D | 0.9159 ± 0.0053 | 0.9079 ± 0.0151 |
| MLP | Real teacher | D+A | **0.8920 ± 0.0055** | **0.8839 ± 0.0050** |
| MLP | Random | D+A | 0.7504 ± 0.0458 | 0.7284 ± 0.0517 |
| MLP | Shuffled teacher | D+A | 0.7617 ± 0.0412 | 0.7398 ± 0.0500 |
| MLP | Random | w/o A | 0.8015 ± 0.0415 | 0.7904 ± 0.0475 |
| MLP | - | w/o D | 0.7903 ± 0.0049 | 0.7775 ± 0.0070 |

Three observations indicate that the teacher outputs contain real and learnable structure despite their chance-level task performance. As direct task predictors, the three label sources are essentially indistinguishable: their AUCs against the true labels on the held-out test set are 0.521 for the real teacher, 0.518 for the random labels, and 0.504 for the shuffled labels (Figure 4B, grey bars). When students are distilled from these sources, however, only the real teacher is reproduced faithfully. At the end of distillation, student-teacher agreement reaches an AUC of 0.986 for MINT and 0.895 for MLP, with per-sample Pearson correlations of 0.965 and 0.775, respectively. Under random or shuffled labels, by contrast, agreement remains near chance (about 0.5), and the student outputs collapse toward an uninformative constant (Figures 4A and 6A). The downstream effects of these seemingly equally uninformative label sources then diverge sharply (Figure 4B). Together, these results suggest that the teacher contains genuine general protein-ligand knowledge that is transferable but not directly aligned with the cofactor-specific objective. This is exactly the setting in which distilling a generalist model into a task-specific specialist is useful.

**Figure 6.**
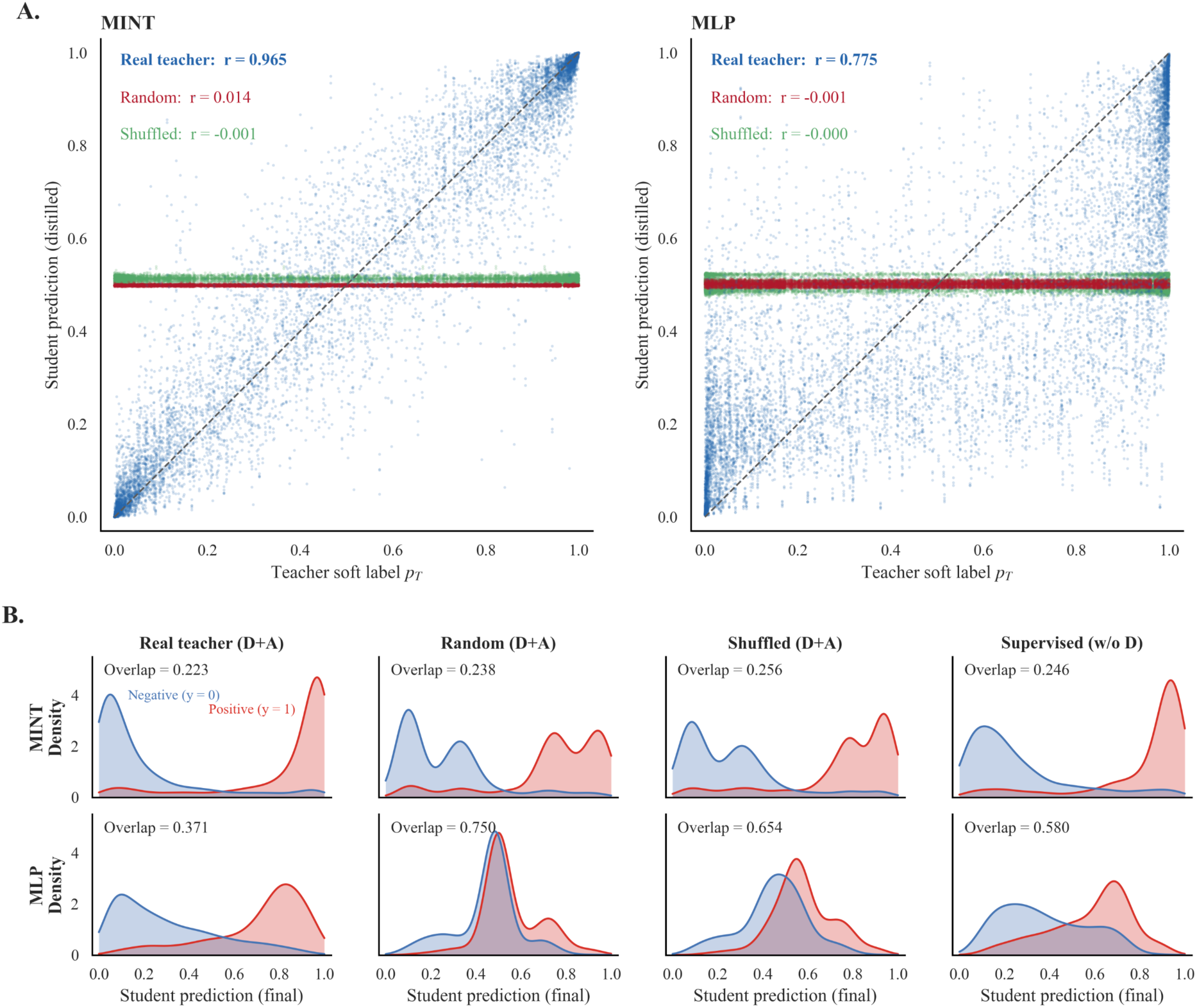
Sample-level effects of distillation and annealing. (A) Per-sample agreement between teacher soft labels and student predictions at the end of distillation (10,000 held-out predictions for each label source: five models × 2,000 enzyme-cofactor pairs; dashed line, identity). Students faithfully reproduce the real teacher’s per-sample scores (blue; Pearson *r*=0.965 for MINT, 0.775 for MLP), whereas under structureless pseudo-labels, random (red) or shuffled (green), the outputs collapse toward an uninformative constant (*r*≈ 0). (B) Class-conditional densities of final student predictions (red, positive; blue, negative) for both backbones under four training configurations, with the overlap coefficient annotated (; 0 = perfect separation, 1 = complete overlap).

Both components of the training strategy contribute to both backbones (Table 3, Figure 5). Removing distillation reduces AUC by 0.016 on MINT (0.932 → 0.916) and by 0.102 on MLP (0.892 → 0.790). Removing annealing while retaining distillation reduces AUC by 0.032 on MINT and by 0.022 on MLP. Each component is therefore beneficial on two architecturally distinct backbones, and in both cases the effect is larger than the fold-to-fold variability of the corresponding models.

The pseudo-teacher controls further show that these gains come from the information carried by the teacher signal rather than from the distillation procedure itself. Replacing the soft labels with random or shuffled values removed the improvement on both backbones. On MINT, the pseudo-label configurations performed at about the level of the supervised reference (0.914 and 0.907 versus 0.916), whereas on MLP they performed below it on average (0.750 and 0.762 versus 0.790), although with substantial fold-to-fold variability (per-configuration SD about 0.04-0.05) (Table 4). The shuffled control is especially informative because it preserves the exact marginal distribution of the teacher scores. The loss of any improvement must therefore result from breaking the sample-wise correspondence rather than from a change in score distribution. This is consistent with a two-stage interpretation: students distilled from the real teacher first learn to reproduce the teacher outputs (Figure 6A) and, with them, the teacher’s own task-level behavior (test AUC at the end of distillation: 0.523 for MINT and 0.518 for MLP). Discriminative performance is then established during the subsequent annealed supervised refinement.

The annealing schedule also acts as a safeguard against low-quality supervision, although its effect depends on the robustness of the overall model design. On MINT, annealing improves performance under every label source, including both the real teacher (+0.032) and random labels (+0.012), and the final class-conditional prediction distributions remain well separated in all configurations (overlap coefficient 0.223-0.256, Figure 6B). The real-teacher configuration achieves the smallest overlap of all, even lower than its supervised reference. On MLP, annealing improves performance only when the real teacher is used (+0.022). Under random labels, annealing does not recover performance (0.750 with annealing versus 0.802 without), and the latter was close to the supervised reference (0.790); the final prediction distributions also deteriorated toward nearly indistinguishable classes (overlap up to 0.750). Consistent with this pattern, MINT varies within a narrow performance range across all six configurations (AUC range 0.032; per-configuration SD≤0.011), whereas the MLP baseline is much more sensitive to the quality of the distillation signal (range 0.142). Annealing therefore gradually downweights supervision that is uninformative or poorly aligned with the target task. This mechanism is effective within a robust architecture, but it is not a universal fix.

### Robustness under family-stratified evaluation

Standard cross-validation on exhaustively paired data can give overly optimistic estimates, and the balanced held-out test set described above includes only 2,000 pairs at an artificial 1:1 positive rate. It therefore does not measure performance at the natural positive rate (0.41%). We therefore repeated the evaluation using an independent family-stratified five-fold split.

Within each family-by-label group, samples were divided into five parts that rotated through training and testing, so that each of the 1,036,680 pairs was held out exactly once and performance could be aggregated over the full dataset at its natural positive rate. Stratifying by family kept the family composition of each fold aligned with that of the full dataset and helped stabilize family-level estimates. Each protein-cofactor pair appeared in either training or testing, but never both. The same protein could appear in both partitions through pairs with different cofactors, which follows from the pair-level formulation of the task, while redundancy among homologous proteins was limited during curation (pairwise sequence identity <0.4). Family-level analyses were restricted to the 14 families with more than ten experimentally supported positive pairs, which together account for 99.1% of all positives, because reliable ranking estimates are not possible below this level of support. All hyperparameters were inherited from the standard setting without re-tuning. Results from this split are reported separately from those of the standard cross-validation setting and are not directly comparable to the balanced-test results.

The family-stratified evaluation reproduced all of the main conclusions from the standard setting (Table 5). MINT with the full pipeline again ranked first (pooled AUC = 0.939), and the full D+A pipeline remained the best configuration within each backbone. The relative importance of the two training components again differed between backbones. On MINT, removing annealing had a larger effect than removing distillation (0.035 versus 0.018 AUC), whereas on the MLP baseline the opposite pattern was observed (0.015 versus 0.099), consistent with the baseline’s stronger reliance on the teacher signal. MINT again outperformed the MLP baseline, and the teacher again performed at chance level (0.514). The close agreement between these two independent split designs (overall AUC 0.932 versus 0.939 for the full MINT model) suggests that the main results are not an artifact of a particular data partition.

**Table 5.** Pooled performance under family-stratified five-fold cross-validation.

| Method | AUC | First-pos rank | Median pos pct | EF@1% |
| --- | --- | --- | --- | --- |
| MINT D+A | **0.9386** | **1** | **0.977** | **27.2×** |
| MINT w/o D (supervised) | 0.9208 | 1 | 0.971 | 19.5× |
| MINT w/o A | 0.9039 | 9 | 0.964 | 18.0× |
| MLP D+A | 0.8947 | 102 | 0.957 | 15.1× |
| MLP w/o A | 0.8795 | 102 | 0.949 | 12.9× |
| MLP w/o D (supervised) | 0.7953 | 86 | 0.879 | 7.0× |
| Teacher (PLXFPred) | 0.5135 | 694 | 0.514 | 0.77× |

Family-level analysis (Figure 7) shows that the advantage of the full MINT model is broad rather than driven by a few dominant families. It ranked first in 11 of the 14 evaluated families (mean rank 1.50), including all eight high-abundance families (*n_pos_* ≥ 200), six of which achieved an AUC above 0.92. The three families in which it was not the top-performing method, biotin (BTN), cobalamin (B12), and pyrroloquinoline quinone (PQQ), were all sparsely annotated. The teacher’s behavior is again informative. Its family-level performance was highly heterogeneous, ranging from moderately informative on S-adenosylmethionine (SAM, 0.693), flavin mononucleotide (FMN, 0.679), and flavin adenine dinucleotide (FAD, 0.645) to systematically anti-correlated on nicotinamide adenine dinucleotide (NAD, 0.399), its 2′-phosphorylated counterpart NADP (0.427), B12 (0.399), and BTN (0.412). The teacher’s near-chance global score therefore appears to reflect cancellation between family-specific alignment and misalignment, rather than a uniform lack of knowledge.

**Figure 7.**
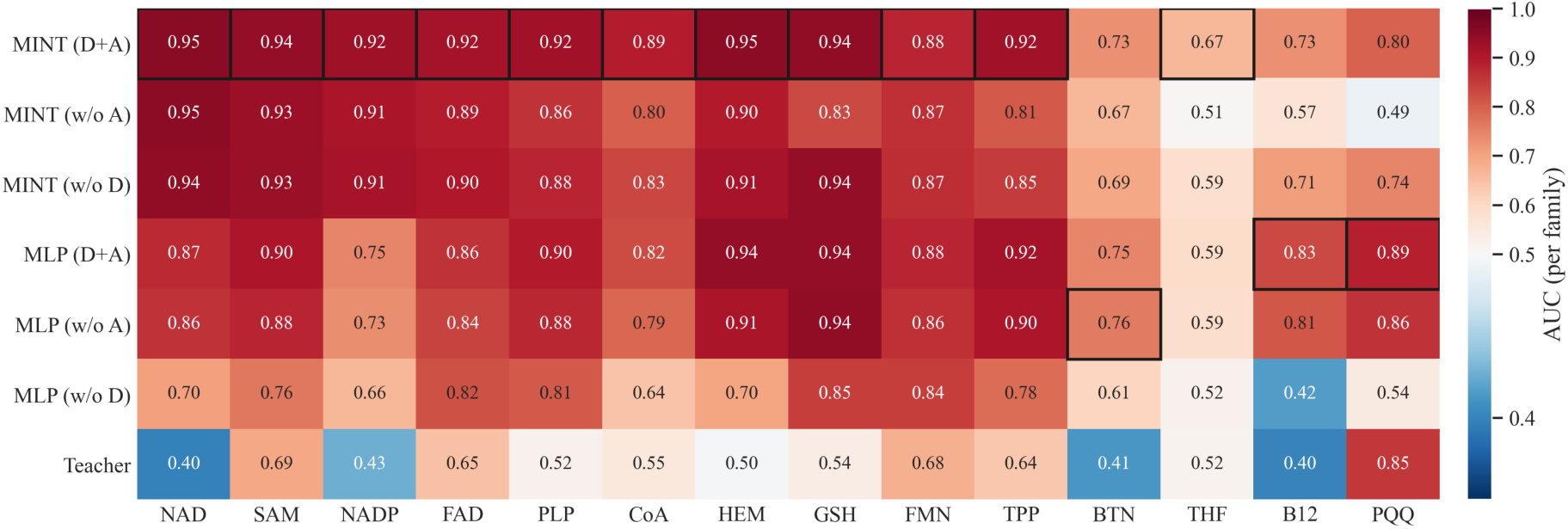
Per-family AUC of the seven methods under family-stratified cross-validation. Fourteen families with more than ten positive samples are shown (columns, ordered by the number of experimentally supported positives); rows correspond to the seven methods. The diverging color scale is centered at 0.5 (chance level), and black frames mark the best-performing method in each family. Family abbreviations are the same as in Figure 1.

The benefit of distillation was greatest where direct supervision was scarcest. Within MINT, the per-family gain of the full pipeline over the supervised variant increased as annotation became sparser: +0.081 for tetrahydrofolate (THF, 36 positives), +0.072 for thiamine pyrophosphate (TPP, 150), and +0.059 for coenzyme A (CoA, 382), compared with +0.008 for NAD (665) and +0.006 for SAM (613). Distillation thus acts as a structured regularizer that provides learning signal where labels are rare, rather than simply amplifying families that are already well represented.

Finally, the pooled family-stratified evaluation approximates a realistic virtual screening setting in which the model ranks about one million candidate pairs at a 0.41% positive rate. In this setting, the full MINT model placed the first true positive at rank 1 and achieved a 27.2-fold enrichment within the top 1% of the ranked list. This was well ahead of the supervised MLP baseline (first true positive at rank 86; 7.0-fold enrichment) and the teacher (rank 694; 0.77-fold).

### Within-protein cofactor ranking

The evaluations above measure pair-level discrimination across the full dataset. A complementary and more application-oriented question is whether a model can rank cofactors for a single protein, which is the setting encountered when annotating an uncharacterized enzyme. To address this, we used the pooled out-of-fold predictions from the family-stratified evaluation to score the 265 candidate cofactors for each of the 3,912 proteins, computed per-protein ranking metrics, and macro-averaged these metrics across all proteins (Table 6).

**Table 6.** Within-protein cofactor ranking under the family-stratified evaluation (macro-averaged over 3,912 proteins).

| Method | Within-protein AUC | Top-1 hit (%) | Top-10 hit (%) | Median best rank |
| --- | --- | --- | --- | --- |
| MINT D+A | **0.938** | **13.6** | **71.3** | **6** |
| MINT w/o D (supervised) | 0.920 | 8.2 | 64.2 | 7 |
| MINT w/o A | 0.913 | 7.7 | 55.4 | 9 |
| MLP D+A | 0.894 | 10.3 | 56.0 | 9 |
| MLP w/o A | 0.876 | 9.3 | 48.5 | 11 |
| MLP w/o D (supervised) | 0.797 | 5.4 | 27.9 | 26 |
| Teacher (PLXFPred) | 0.515 | 0.2 | 5.2 | 140 |

The full MINT model again ranked first on every metric (Table 6). It achieved a within-protein AUC of 0.938; for 71.3% of proteins, a true cofactor appeared within the top 10 of the 265 candidates, and the highest-ranked true cofactor had a median rank of 6 among 265 candidates. Both components of the training strategy contributed, as removing distillation or annealing degraded within-protein ranking on both backbones, and the MLP baseline performed worse than all MINT configurations. In contrast, the teacher performed at chance level (within-protein AUC 0.515). Only 5.2% of proteins had a true cofactor within the top 10, compared with a random expectation of about 4.1%, and no protein had its best cofactor ranked first; the top-1 hit rate was 0.2%, below the expected value of about 0.4%. These results confirm that general interaction scores do not order a protein’s candidate cofactors in a useful way.

The nicotinamide families again show the teacher’s limitation most clearly. Among the 621 proteins annotated to bind NAD but not NADP, the teacher ranked the true cofactor ahead of its 2′-phosphorylated counterpart in only 14.0% of cases, indicating a systematic inversion, whereas the full MINT model did so in 88.1%. In the corresponding group of 546 proteins annotated to bind NADP, the figures were 56.3% for the teacher and 95.8% for MINT. Ranking cofactors within a protein therefore requires exactly the kind of single-phosphate resolution that the general teacher lacks and that distillation with annealed refinement restores.

### Case study: NAD/NADP discrimination

To illustrate how teacher-guided learning resolves a concrete and biologically important discrimination problem, we examined the nicotinamide cofactor families in detail. NAD and NADP differ by a single phosphate group at the 2′ position of the adenosine ribose (Figure 8A), yet this small chemical difference underlies their distinct metabolic roles, and engineering NAD/NADP specificity is a well-established goal in enzyme design. This near identity makes the nicotinamide pair the most difficult discrimination problem in the dataset. The two cofactors are chemically almost indistinguishable and bind evolutionarily related pockets in nearly identical poses, so the predictor must resolve differences at the level of a single phosphate and, structurally, a single pocket residue.

**Figure 8.**
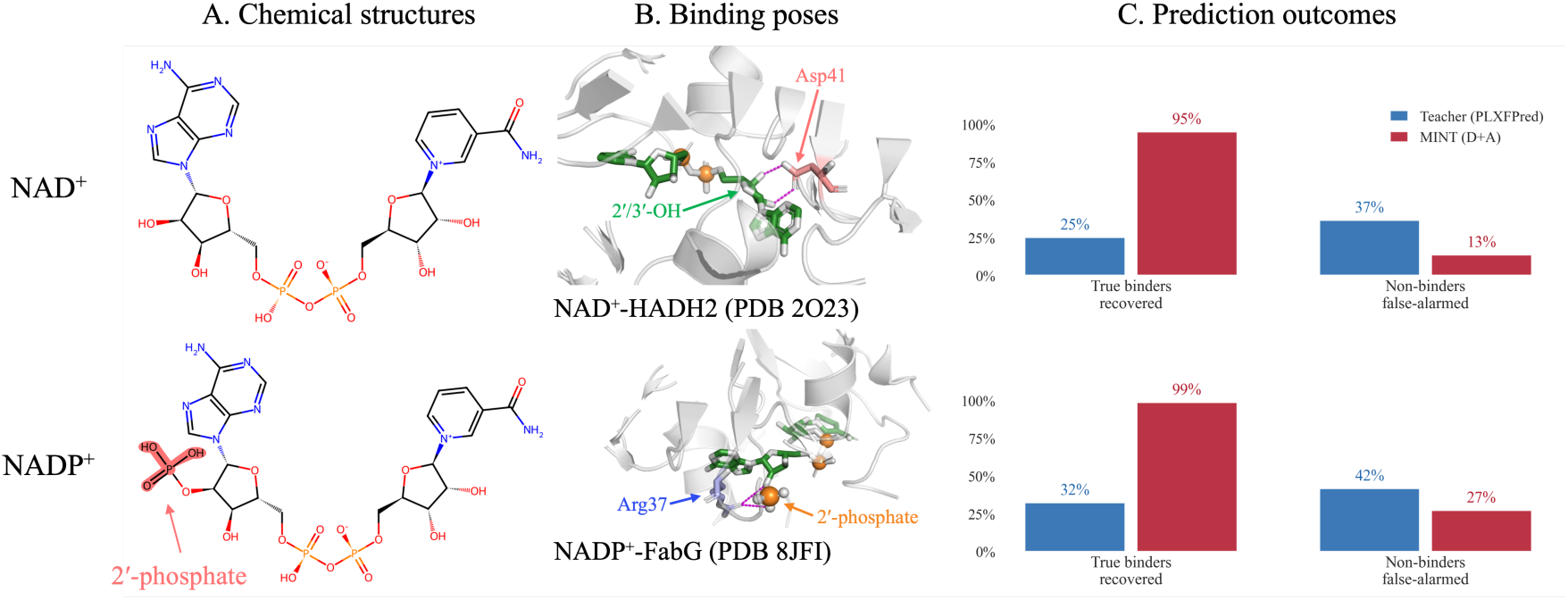
Case study: NAD/NADP discrimination. (A) Two-dimensional structures of NAD^+^ and NADP^+^; the only difference, the 2′-phosphate on the adenosine ribose, is highlighted. (B) Binding poses in two short-chain dehydrogenase/reductase (SDR) complexes. Top: NAD^+^ bound to human HADH2 (PDB: 2O23, chain A); Asp41 (salmon) forms hydrogen bonds with the ribose 2′/3′-hydroxyl groups (2.6 Å). Bottom: NADP^+^ bound to FabG from Helicobacter pylori (PDB: 8JFI, chain A); Arg37 (blue) anchors the 2′-phosphate (3.3-3.5 Å). Cofactors are shown as green sticks, phosphorus atoms as orange spheres, and hydrogen bonds as magenta dashed lines; the two chain-A protomers were superposed in PyMOL (super; backbone RMSD 0.94 Å over 159 C*α* atoms). (C) Recovery of true binders and false-alarm rate for non-binders at the default operating point (score >0.5), averaged across the five cross-validation models on the held-out test set (NAD: 179 positives / 104 negatives; NADP: 130 / 36).

The teacher fails at exactly this level of resolution. Under the family-stratified evaluation, its family-level AUC falls below 0.5 for both nicotinamide families (NAD: 0.399; NADP: 0.427). At its default operating point (score >0.5), this failure becomes even clearer: on the held-out test set, it recovers only 25.1% of the 179 true NAD binders and 32.3% of the 130 true NADP binders, while incorrectly labeling 36.5% (NAD) and 41.7% (NADP) of non-binders as positive. In other words, its false-alarm rate exceeds its recovery rate, indicating that its scores on these families provide little useful signal (Figure 8C). A general protein-ligand interaction model trained across many ligand classes does not retain the single-phosphate resolution required for cofactor preference prediction. This limitation motivates the need for a dedicated cofactor-preference predictor: in this domain, the biologically decisive distinctions fall below the resolution of general-purpose models. After distillation and annealed refinement, MINT recovers exactly this level of resolution. Averaged across the five cross-validation models, it recovers 95.0% (NAD) and 98.6% (NADP) of true binders on the held-out test set, with false-alarm rates of 13.5% and 27.2%, respectively, and achieves family-stratified AUCs of 0.952 (NAD) and 0.923 (NADP).

Structural evidence helps explain what the teacher misses. In the short-chain dehydrogenase/reductase (SDR) superfamily, cofactor preference is determined by a single pocket position: NADP^+^-dependent enzymes place a positively charged residue there to anchor the 2′-phosphate, whereas NAD^+^-dependent enzymes place an acidic residue that would repel it. In the NADP^+^ complex of FabG from Helicobacter pylori (PDB: 8JFI, 2.38 Å), the guanidinium group of Arg37 forms direct hydrogen bonds with the distinguishing 2′-phosphate (3.3-3.5 Å; Figure 8B, bottom) [37]. In the NAD^+^ complex of human HADH2 (PDB: 2O23, 1.20 Å), the equivalent position is occupied by Asp41, which instead forms two hydrogen bonds with the ribose 2′- and 3′-hydroxyl groups (2.6 Å; Figure 8B, top), an arrangement that is electrostatically incompatible with a 2′-phosphate [38]. The two proteins are structural neighbors (backbone RMSD 0.94 Å over 159 Cα atoms), and the two cofactors adopt nearly identical extended poses. Their preference is effectively determined by this single-residue, single-phosphate exchange. Consistent with having learned the cofactor-specific objective, the three 8JFI-NADP positive pairs in the held-out test set received a median score of 0.969 across the five models, precisely in the regime where the teacher’s ranking is least reliable. This case study closes the loop: knowledge distilled from a general interaction model, once realigned to the cofactor-specific objective, recovers the fine-grained discrimination between chemically near-identical cofactors that is required for enzyme cofactor-specificity engineering and orphan-enzyme annotation.

## DISCUSSION

This study shows that protein-cofactor selectivity prediction can benefit substantially from transfer learning under limited supervision. The proposed MINT framework outperformed the MLP baseline, its ablated variants, and the teacher model under two independent cross-validation designs, with nearly identical overall performance (AUC 0.932 versus 0.939) and the same ranking of methods in both settings. These results indicate that combining knowledge distillation, progressive annealed adaptation, and multimodal representation learning is effective for this task, and that the main conclusions are robust to the choice of data split.

An important finding is that the value of the teacher model lies mainly in the supervision it provides during training rather than in its direct downstream performance. Although PLXFPred performed near chance when applied directly to cofactor specificity prediction, students reproduced its soft outputs with high fidelity and improved substantially after training on them, whereas label sources with similar task-level quality but no structured content, such as random or shuffled labels, provided no such benefit. Task alignment and information content therefore appear to be orthogonal properties of a teacher model: a model trained on a broader interaction task can encode transferable structure even when its native predictions are not aligned with the downstream objective. The family-level analysis sharpened this point further, showing that the teacher’s near-chance global performance masks strong but opposing family-specific effects, ranging from moderately informative behavior (SAM) to systematic anti-correlation (NAD/NADP). This decoupling provides direct support for distilling general protein-ligand models into task-specific specialists, rather than either applying them directly or dismissing them as uninformative.

The ablation results further show that the benefit of teacher-guided learning depends on the interaction between the training strategy and the student architecture. Both distillation and annealing contributed on two independent backbones, although their relative importance depended on the backbone: annealing had the larger effect for MINT, whereas distillation contributed more for the simpler MLP baseline. The two backbones also differed markedly in robustness. MINT maintained stable performance and well-separated prediction distributions under every supervision regime tested, including deliberately corrupted teacher signals, whereas the simpler MLP baseline benefited substantially from the real teacher but was destabilized by low-quality supervision. Annealing emerged as an important safeguard. By gradually shifting the optimization objective from teacher guidance toward hard-label supervision, it allows the model to retain useful transferred structure while filtering out signals that are uninformative or misaligned with the target task, including the teacher’s own task-level biases. That this filtering operated reliably in MINT but not in the less robust baseline suggests that gradual task transition and a strong student design are complementary rather than individually sufficient.

The family-level analysis also suggests a practical rule of thumb: the relative benefit of distillation increases as direct annotation becomes scarcer. This pattern is consistent with the view of teacher guidance as a structured regularizer that constrains learning when labels are limited, and it suggests that the framework provides its greatest marginal value precisely in the low-resource setting that motivates its use.

The results also have direct practical implications. At the natural positive rate of the dataset (0.41%), the full model ranked the first true positive at the top of a candidate list of about one million pairs and achieved 27-fold enrichment within the top 1% of ranked candidates. It also recovered the true cofactors of individual proteins from each protein’s own candidate list, supporting applications such as prioritizing candidate cofactors for uncharacterized enzymes. The nicotinamide case study illustrates the level of resolution that matters in enzyme engineering: the model distinguishes NAD from NADP preferences, even though these cofactors differ by only a single phosphate group and the teacher is systematically anti-correlated on these families. This kind of discrimination is directly relevant to cofactor-specificity engineering workflows and suggests that the framework could complement experimental engineering efforts. At the same time, both evaluation designs remain within an interpolation setting, in which test proteins and cofactors are drawn from the same curated space as the training data. Generalization to proteins that are entirely unrelated to the training set remains to be established.

Several limitations should be noted. Experimentally supported positive samples remain scarce, and the negative set was defined largely by the absence of evidence, which may introduce label noise. The current task formulation is limited to pairwise prediction and does not explicitly model the broader biochemical context. In addition, the individual contributions of the different input modalities were not fully disentangled. Future work should therefore examine more reliable strategies for negative-sample construction, modality-specific ablations, and validation on independent external datasets.

Overall, these results support viewing protein-cofactor selectivity prediction as a transfer learning problem rather than a purely task-specific supervised classification problem. MINT provides an effective framework for improving prediction under sparse annotation, and the combination of distillation with annealed adaptation offers a useful strategy for related low-resource molecular recognition tasks.

## CONCLUSION

In conclusion, this study presents MINT, a multimodal teacher-supervised framework for protein-cofactor selectivity prediction. By combining knowledge distillation from a general protein-ligand interaction model with progressive annealed adaptation and a multimodal student architecture, MINT achieved the best overall performance among all methods evaluated. This advantage was consistent across two independent evaluation designs: a balanced held-out test set (AUC = 0.932) and a family-stratified split at the natural positive rate (pooled AUC = 0.939). Mechanistically, we show that the teacher’s value lies in the transferable structure it provides rather than in its own task accuracy. Students reproduced the teacher’s outputs with high fidelity, whereas structurally uninformative pseudo-labels did not provide the same benefit, and annealing progressively realigned the transferred knowledge to the cofactor-specific objective. The NAD/NADP case study further showed that the resulting model can resolve single-phosphate-level differences that the general teacher cannot, illustrating the level of resolution required for enzyme cofactor-specificity engineering. Taken together, these findings show that effective cofactor prediction depends not only on the downstream objective itself, but also on the training path used to reach it. The proposed framework provides a practical solution for cofactor specificity prediction under limited annotation and may also be useful for other low-resource biochemical prediction tasks.

## AUTHOR INFORMATION

### Author Contributions

Y.Y.: investigation, methodology, software, data curation, formal analysis, visualization, writing - original draft. X.L.: methodology, formal analysis, visualization. H.D., Y.A.: writing - original draft, writing - review and editing. J.G.: conceptualization, supervision, writing - review and editing. All authors have read and agreed to the published version of the manuscript.

### Data Availability

The implementation of the proposed method will be made publicly available on GitHub at https://github.com/AIDD-YE/MINT.

## Acknowledgements

This research was supported by the Science and Technology Development Fund (FDCT) of Macau (0004/2025/RIA1) and Macao Polytechnic University (RP/CAI-01/2023).

